# Single–molecule biodosimetry

**DOI:** 10.1101/2024.12.05.627019

**Authors:** Michael Lamontagne, Shannon M. Newell, Ileana Pazos, Ronald Tosh, Jerimy Polf, Michael Zwolak, Joseph W. F. Robertson

## Abstract

Inferring characteristics of radiation exposure using biological molecules is extremely challenging. Current methods, in particular, lack a clear connection between dose and molecular response. Here, we demonstrate that resistive–pulse nanopore sensors enable single–molecule biodosimetry by quantifying the frequency of double–strand DNA scissions versus gamma radiation dose. The resulting response curve shows an elongated Gaussian behavior, reminiscent of cell survival rates versus dose. We demonstrate that the competition of radical damage of DNA—i.e., single–strand lesions that lead to breakage—with bimolecular radical loss captures the form of the response. Our sensors and protocol provide a foundation for numerous technological advances. These include rapid dosimetry for triage in emergency situations and *ex vivo* monitoring of radiotherapy effectiveness in order to tailor treatment to patient– and tumor–specific response.

---

Understanding the biological consequences and damage mechanisms of ionizing radiation is central to cancer therapy [1]. Quantifying radiation exposure, in particular, enables post–therapy assessment of the delivered dose [2] and will be important in other scenarios, such as radiological accidents [3] or nuclear conflicts [4]. Moreover, beyond the acute effects of high–dose exposure (*>* 2 Gy), the accumulated impact of low–dose exposure (≈ 100 *µ*Gy), such as from medical imaging or working in mines, is a poorly understood factor in assessing health [5]. Current tools are often inadequate for these medical and emergency applications [6]. For instance, the “gold standard” for retrospective dosimetry—dicentric chromosomal analysis—requires *>* 48 h of preparation time for cell culturing after sample collection to produce actionable data [7]. Other chromosomal counting assays and proteomic tools have similar issues. Leveraging advances in biotechnology and nanoscience is a promising route to overcome such challenges and provide tools to quantitatively assess damage and mechanisms, as well as to rapidly and accurately measure the exposure of individuals under a broad range of conditions [5].

In this regard, the most well–studied biomarker for assessing radiation damage is DNA. It has long been known that radiation–induced damage to genomic DNA can lead to cell death [8, 9], with as little as one double– strand break (DSB) being sufficient in certain circumstances [10]. In living systems, cell death can be mitigated by the action of DNA repair enzymes [11, 12]. However, in a buffer containing DNA, in addition to the lack of molecular packing (as in cells) [13], radiation damage proceeds unperturbed by repair enzymes. This yields a more straightforward picture of the damage mechanisms and rates. For DNA, damage includes both *direct* and *indirect* lesions on the bases or sugars. In practice, irradiation deposits the most energy in water—the most abundant species by mass. This initiates a cascade of reactive hydrolysis products, including 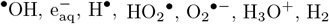, and H_2_O_2_ [14]. Of these,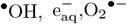 and H_2_O_2_ are particularly damaging to DNA, with ^*•*^OH specifically associated with strand scission [15– 17]. H-abstraction on the ribose occurs in 10% and 20% of all ^*•*^OH reactions with DNA, with *≈* 20% of these abstraction reactions leading to strand–break reactions [18, 19].

Nanopore sensors emerged as a tool to study the properties of DNA and other biopolymers in 1996 [20]. The core sensing principle is similar to other biophysical research tools, such as the Coulter counter [21, 22]: A thin membrane (with a narrow pore) partitions an electrolyte solution, where a voltage drop drives an ionic current from one side to the other. When the analyte is driven through the pore—via fluid pressure for a Coulter counter or electrophoretically for DNA—a resistive pulse occurs in the current. The capture process itself is complex. The electric fields and polymer dynamics [23, 24], as well as the interplay between electrophoretic and electroosmotic forces with the effective charge of the molecule [25, 26], all play a role. The molecular volumes of nanopores (rather than cellular volumes of Coulter counters) allow the system to measure single molecules. In particular, the magnitude of the resistive pulse is proportional to the volume of the pore occupied by the translocating molecule [27, 28]. For long, uniformly–charged molecules, such as DNA, integrating the pulse yields a value proportional to the molecule’s length [29, 30].

Since their introduction in 2001 [31], solid–state nanopores have become a workhorse of single– biomolecule sensing [32]. The most common approach is to fabricate nanopores in ultra–thin silicon nitride membranes, which enables sampling above megahertz frequencies and therefore maximizes discrimination of sub-level blockades within individual translocations [33, 34]. Yet, these pores are highly susceptible to fouling. Other geometries and materials, such as conical pores in quartz glasses, are similarly effective for sensing [35, 36]. On– demand laser–pulled capillaries, for instance, provide a simple, deployable alternative to more difficult to fabricate pores [37, 38]. Yet, like other solid–state pores, they have highly variable pore characteristics (surfaces, taper, etc.). Despite this, we employ pulled quartz nanopipettes to demonstrate single–molecule biodosimetry. We will provide a method that can tolerate large pore–to–pore variation and enable precise, quantitative measurement of DNA damage, while being simple to use.

## I. RESULTS

The single–molecule biodosimetry approach we develop employs nanopores to quantify double–strand breaks from an irradiated sample. Here, we expose a series of aqueous solutions containing 2.5 kbp DNA, Fig. 1a, to gamma radiation with doses up to 15 Gy, which results in double–strand breaks, Fig. 1b. In order to extract quantitative information, two longer DNA molecules (here, 5 kbp and 10 kbp) are added post– irradiation as simultaneous concentration and molecular– length standards.

**FIG 1.**
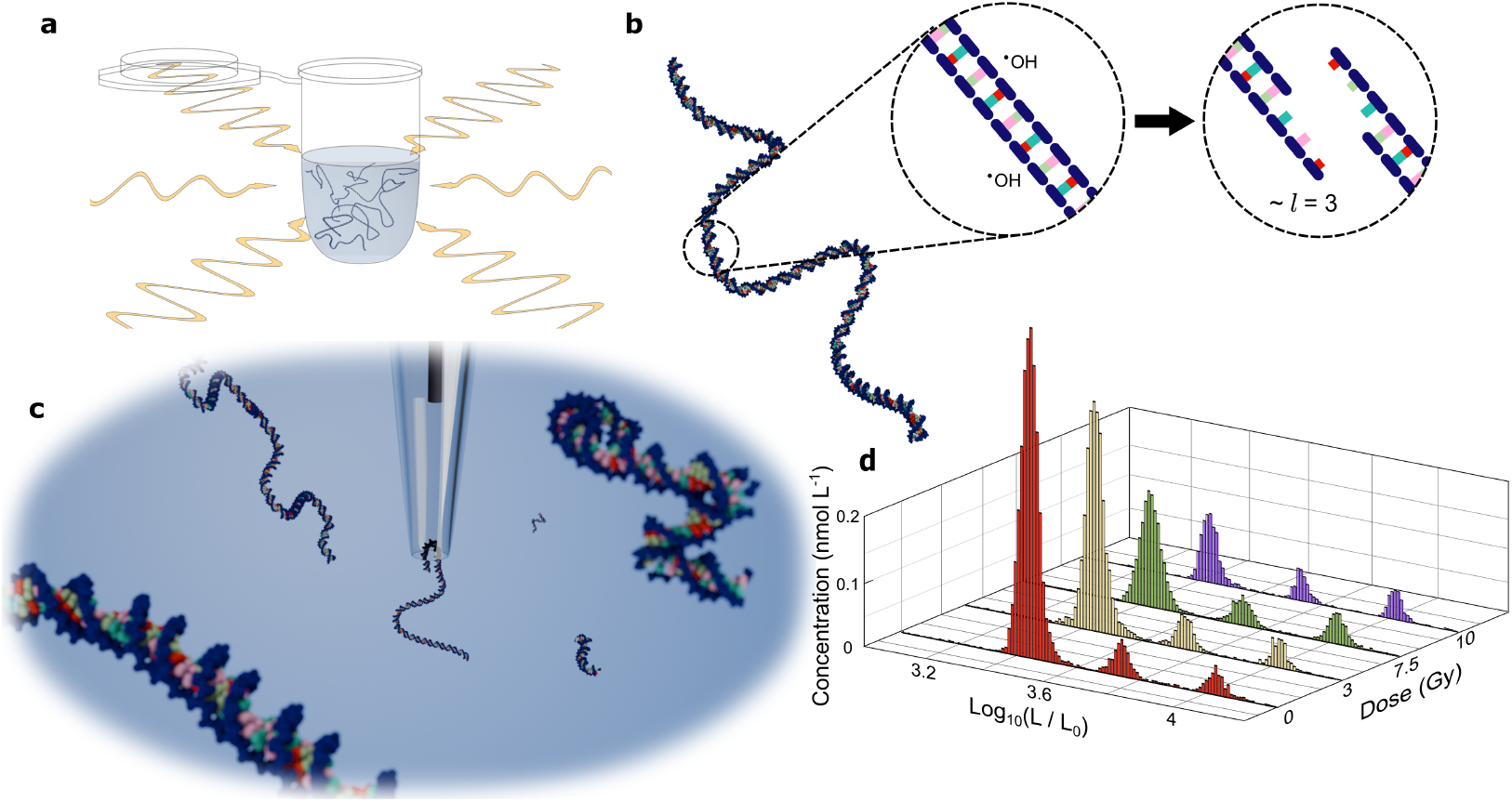
Nanopore–based single–molecule dosimetry. **a**, An aqueous solution of 2.5 kbp DNA is irradiated with a ^60^Co calibrated gamma source. **b**, Single–strand breaks in the sugar–phosphate backbone accumulate until two breaks occur close enough (here shown as *l* = 3 bases) to compromise the stability of the molecule and cause a double–strand break. **c**, Irradiated DNA sample is quantified post–exposure with a glass nanopipette. **d**, Histograms of the concentrations of the irradiated 2.5 kbp and unirradiated internal standards of 5 kbp and 10 kbp DNA as a function of dose (*L*_0_ = 1 bp). Absolute size and concentration are calculated from the ECD and capture frequencies by calibrating against these two internal standards.

During measurement, DNA is electrophoretically driven into the mouth of a glass nanopipette, Fig. 1c. The resulting current blockades—the resistive pulses—are analyzed and compiled into a histogram of molecular length, Fig. 1d. The equivalent charge deficit (ECD) [29]—the total area of a resistive pulse (shaded in Fig. 2a,b)—yields information on the molecular size for a single nanopore.

**FIG 2.**
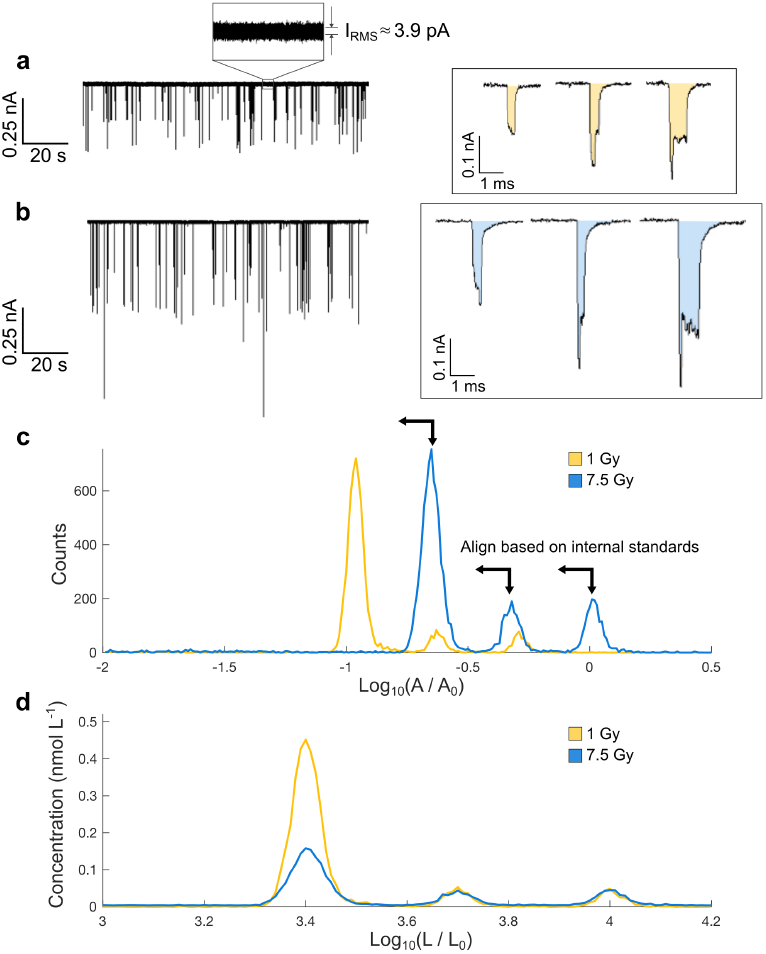
Molecular standards as an internal calibration and ruler. **a**, Ionic current versus time showing resistive pulses of the 5 kbp and 10 kbp DNA internal standards and 1.0 Gy gamma irradiated 2.5 kbp DNA. **b**, Ionic current of an identical DNA mixture in which the 2.5 kbp DNA has been irradiated at 15.0 Gy. Characteristic current events for the irradiated length, 2.5 kbp (left), and the two internal standards, 5 kbp (center) and 10 kbp (right), are shown for 1.0 Gy and 7.5 Gy enlarged in the boxes. The ECD is the area, A, shaded in orange or blue. **c**, Without calibration, by visual inspection, the 5 kbp 1.0 Gy events are closer in magnitude to the 2.5 kbp events in the 7.5 Gy run than are to the corresponding 5 kbp events (here, *A*_0_ = 1 pC). **d**, This is corrected by aligning the internal standard peaks horizontally (here, *L*_0_ = 1 bp). The concentration is calculated by comparing the average integral of the 5 kbp and 10 kbp peaks, to the known concentration at which they were added. After alignment, the final 1.0 Gy and 7.5 Gy histograms show a decreasing concentration of intact 2.5 kbp DNA with dose. The integrated area of the 2.5 kbp peak is taken as its measured concentration for each dose.

Yet, it is not feasible to use the same pore for every dose, as there can be both cross contamination of samples and pore fouling (typically after less than two hours, depending on analyte concentration and pore size). These are common limitations for solid–state nanopores and we must calibrate pore-to-pore variations between a series of pores prepared identically. Indeed, as seen in Fig. 2c, there is a large variability in the uncalibrated ECD peaks for each dose. Building on prior work that calibrated the concentration using the capture frequency of an internal standard [39], we introduce two internal standards, which provides a correction for *both* size and concentration simultaneously, see Fig. 2c,d. Moreover, it also yields quantitative data about uncertainty.

The resulting dose–response curve agrees well with a model of double–strand breaks originating from two nearby single–strand lesions. Regardless of whether those lesions form directly from radiation damage or indirectly from reaction of the DNA with radicals, they deplete the initial intact DNA concentration Φ_0_ according to a universal form (see the Methods) in the small *P*_Λ_ limit,

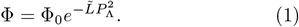

Here, 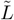 is the effective length (in number o f base pairs) of the DNA and *P*_Λ_ is the lesion probability, i.e., the likelihood of individual single–strand lesions. The effective length, 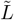, is proportional to the actual length of DNA and, unlike *P*_Λ_, independent of dose. While Eq. (1) is general, we consider only indirect damage via radical reactions, as the mass fraction of DNA is very small.

To lowest order in *D*, the probability to generate a lesion should grow linearly with the energy deposited per unit mass, or dose, i.e., *P*_Λ_ *∼ D*. A reaction model where radicals damage nucleotides and are also subject to (effective) unimolecular decay gives such a linear dependence provided the initial radical concentration formed by water radiolysis itself is linearly dependent on *D*. Figure 3 suggests Gaussian decay of Φ with *D*. Yet, a low–dose (*≤* 3 Gy) fit (the red, dotted line in Fig. 3), where this model should be valid, gives significantly less intact DNA than expected at large *D*. This indicates that another important reaction or other process must be present.

**FIG 3.**
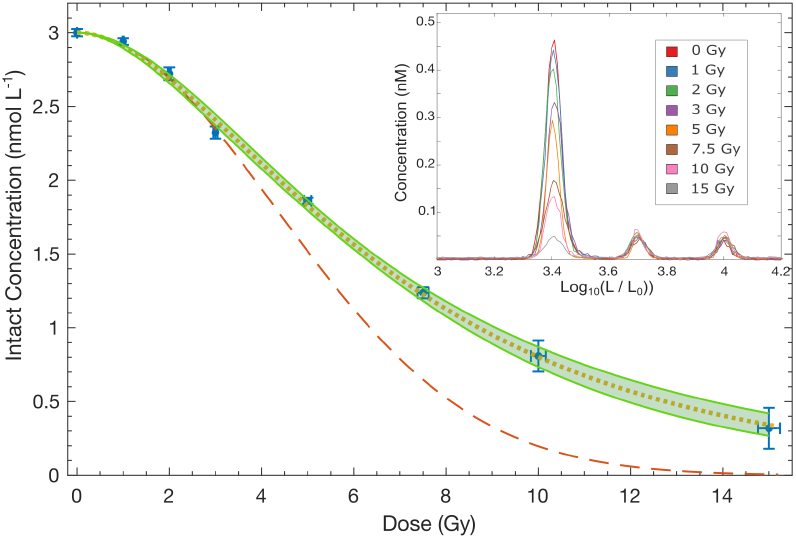
DNA dose response. The intact DNA concentration (blue circles and error bars) versus dose shows that the response is roughly Gaussian. At very low dose, one expects such behavior since bimolecular radical decay will be negligible. However, the best fit (red, dashed line) at low dose,*≤* 2 Gy, to a Gaussian model does not work well at high doses, overestimating the loss of intact DNA. A best fit to the whole range of data (not shown) similarly does not fit well (overestimating intact DNA at intermediate doses and underestimating at high doses), as the shape of the curve is not actually Gaussian in *D* but an elongated Gaussian. Including bimolecular decay yields a two parameter model and a high–quality fit to the data (orange, dotted line). The inset shows the measured concentration of three key molecules. The concentration of 2.5 kbp DNA decreases with increasing dose. Error bars are a scaled average of the standard deviation between the measured concentrations of the 5 kbp and 10 kbp internal standards (that were fixed between runs). The scaling is based on the length of the dataset and the number of 2.5 kbp events recorded at a particular dose.

Bimolecular decay of radicals, i.e., two radicals combine to form an inert species, may be such a reaction[14]. If this loss mechanism is present, the intact DNA still follows Eq. (1) but with a defect probability

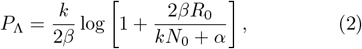

where *k* is the rate constant for DNA damage, *β* the bi-molecular radical decay rate, *α* the unimolecular radical decay rate, *N*_0_ the initial nucleotide concentration during irradiation, and *R*_0_ = *GD* is the initial radical concentration. In the latter, *G*, a constant of proportionality relating *R*_0_ to dose, *D*, is the radiation chemical yield [40]. This model has several parameters. Yet, as a fit to experimental data, it has only two effective parameters,

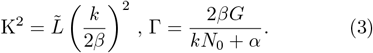

With K and Γ, the experimental data is fit extraordinarily well (green, dashed line in Fig. 3).

We note that these two parameters are correlated, with a decrease in one generally occurring with an increase in the other (i.e., KΓ is roughly constant since the initial decay is Gaussian). The best fit to the data (without accounting for uncertainty) is K = 1.23 and Γ = 0.155 Gy^*−*1^. Including uncertainty, the fit is in Fig. 3, with the main line the average intact concentration from the joint distribution of K and Γ, and the shaded region given by plus and minus one standard deviation. Treating the two parameters independently gives K = 1.3 *±* 0.4 and Γ = (0.16 *±* 0.05) Gy^*−*1^, yet this does not capture their correlations.

Extraction of any of the fundamental parameters in Eq. (3) (i.e., 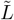*k,α,β* or *G*) requires either tuning various quantities (such as concentrations) or independently measuring or estimating some subset of those parameters. Taking 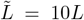 (see the Methods) yields *k/β* = 0.017 *±* 0.005. Further, assuming that the radical scavenging rate, *α*, is of the order of the nucleotide damage rate (e.g., that a dominant decay route is to create DNA damage that does not lead to breakage and thus *α ≈ kN*_0_), we find *G* = (242 *±* 21) nmol·L^*−*1^Gy^*−*1^ or, in conventional units, *G* = (2.3 *±* 0.2) radiolysis species per 100 eV deposited. This value is consistent with prior measurements of the *G* value for ^*•*^OH, which give 2.56 to 5.5 molecules per 100 eV [14, 41]. We can instead take the reported *G* values and use that as the input to extract *k/β* and 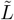. For *G* = 2.56 molecules per 100 eV, *k/β* = 0.019 *±* 0.008 and 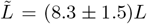 We note 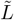 yields *l ≈* 5 bp—the length scale for which SSBs on opposite strands lead to a DSB, which is in line with prior measurements between 2 bp and 16 bp for high and low ionic strengths, respectively [42].

## II. DISCUSSION

Measuring universal, reproducible trends with multiple solid–state nanopores, even pores that are nominally identical, has generally proven elusive. Pore–to– pore variability in diameter, internal taper, and surface charge and roughness, among other factors, has a significant impact on characteristic molecular signatures, such as ECD. We have taken a two–pronged approach to overcome the challenges in getting quantitative trends. Firstly, we employ relatively large nanopipettes (i.e., ≈ 12 nm 17 nm), which leads to a number of advantages over reliance upon smaller pores in addition to ease of use: (i) Large pores frequently enable collection of large data sets (approximately 10 k events before pore fouling); (ii) They are more robust, filling the narrow sensing region of the tip completely a larger percentage of the time and giving a higher yield of pores with very low baseline noise (*i*_*RMS*_ ≈ 4.4 pA); and (iii) They have a flat capture probability across the size range of the three target molecules as shown in Fig. 4c. Incidentally, the nanopipettes also enable recovering the sample, as opposed to traditional solid–state pores where one would have to pipette out the sample and risk contamination. Secondly, we correct for the analyte size and capture–rate fluctuations that result from the natural pore–to–pore variability by using the fixed sizes and concentrations of the two internal molecular standards, which allows us to infer the concentration of all molecules in our target size range, from 2.5 kbp to 10 kbp, for our nanopipettes.

**FIG 4.**
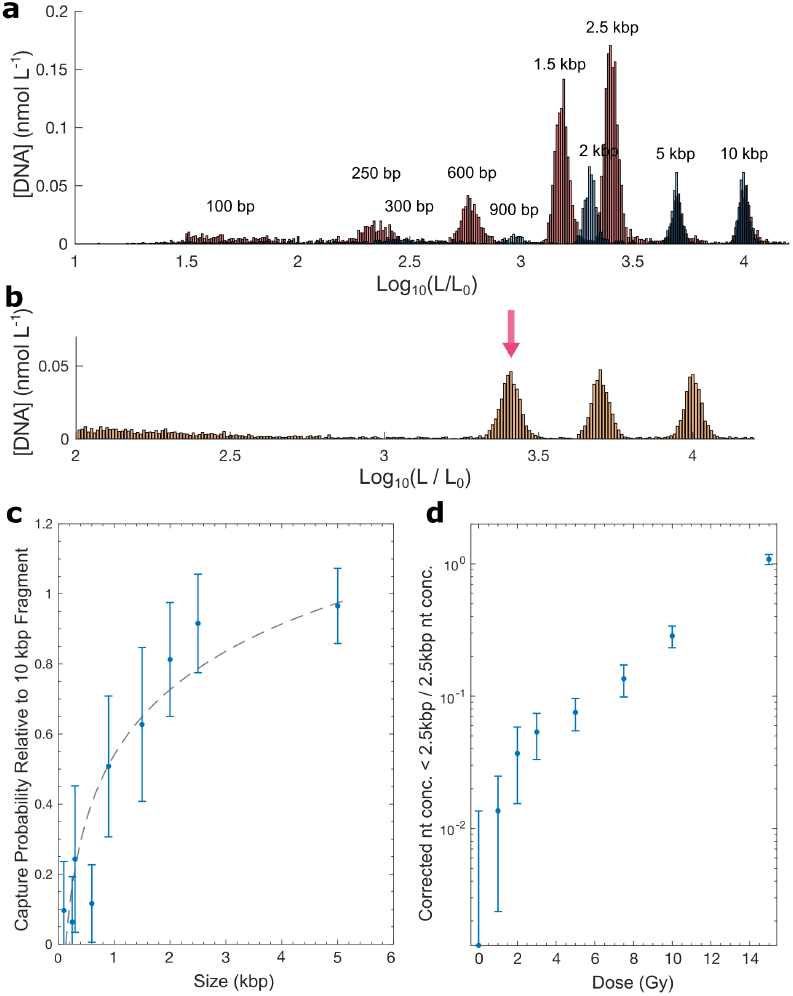
Fragment concentration bias. **a**, A comprehensive molecular ladder of ≤ 10 kbp DNA run through a pipette prepared identically to those used in the primary experiments. Separate ladders containing 100 bp, 250 bp, 600 bp, 1.5 kbp, 2.5 kbp, 5 kbp, and 10 kbp fragments (red), and 300 bp, 900 bp, 2 kbp, 5 kbp, and 10 kbp fragments (blue) were recorded and aligned as described in the Methods to minimize overlap. **b**, Molecular size distribution for all captured molecules after exposure to 15.0 Gy of radiation. Fragments between 100 bp and 2.5 kbp are visible below the 2.5 kbp peak indicated with the red arrow (*L*_0_ = 1 bp). **c**, The probability, *P*_*C*_, of successfully capturing a DNA fragment of a given size as calculated from the ladder shown in **a**. For nanopipettes of this size, *P*_*C*_ is relatively flat in the 2 to 10 kbp range, and decreases logarithmically for fragments *≤* 2 kbp in size. Error bars are the standard deviation of the best–fit parameters of a logarithmic fit (black, dotted line) to the capture rates of the ladder run through three identically prepared pipettes. **d**, The ratio of nucleotides in fragments (the difference between a “fragment” and the intact Gaussian fitted peak being *>* 2 standard deviations) to nucleotides in the intact 2.5 kbp DNA after correcting for the deflated *P*_*C*_ of small molecules. After correction, the fragment-to-intact ratio is monotonically increasing with dose.

This approach enables low uncertainty, molecule–by– molecule measurement of the intact DNA concentration. Yet, using larger pores and optimizing the standards around them is not without trade off: We lose resolution of the concentration of small fragments. Figure 4a shows the concentration across a wide range of molecular sizes, where the weight on the smaller fragments is smaller than expected. There are two primary reasons for this. For small molecules, it is not possible to fully distinguish capture and translocation, partial capture, and system noise. Also, there is a complicated interplay of physical processes on the capture of molecules diffusing near the terminal aperture. Generally speaking, for negatively charged surfaces, the electro–osmotic force on the exterior of the tip of the nanopipette, mediated through fluidic drag interactions, creates a spherical region of pre– concentrated DNA around the pore mouth. Competition occurs between many molecules fluctuating in and out of the high electric field immediately inside the vestibule of the nanopipette, ultimately giving single–file capture. Overall, this latter factor dominates and leads to a lower number of small molecule (relative to the pore) translocations than their concentrations would indicate [23, 25].

In other words, the correction of capture events needs to be *non–linear*. DNA standards with a size of the same order as our target DNA permits a linear correction for the larger size molecules but under weights small fragment events. As a preliminary estimate, we measure the bias of a ladder of known fragment concentrations below the intact DNA size (shown in Fig. 4c.) This is done with three nanopipettes identically prepared to those in the primary experiments. We note that this provides an average correction factor rather than a pore–by–pore calibration as was done for the primary experiments. Consequently, the error bars are larger for the small fragments. Despite this lack of optimization, the corrected ratio of measured DNA fragments (in nucleotides) to intact 2.5 kbp DNA (in nucleotides) monotonically increases with dose, see Fig. 4d. In some scenarios, such as emergency exposure, the initial concentration of undamaged DNA in a sample will not be known. Under such conditions, a ratiometric approach—i.e., measuring the concentration of fragments relative to intact DNA— may enable determination of the dose to the needed accuracy by employing ratio–dose response data like that in Fig. 4d. To apply the ratiometric approach rigorously, one would first need to overcome many deficits in the theory of nanopore capture physics. We anticipate that ratiometric data can be significantly improved using pores optimized for quantitative measurement of molecules on the order of the fragment size.

## III. CONCLUSION

We demonstrate single–molecule biodosimetry using quartz nanopipettes. Our results give the first quantitative, direct connection between dose and DNA response. Calibration against internal molecular standards, as we introduced here as a parallel to DNA electrophoresis ladders, is essential for the acquisition of fully quantitative results and will be critical to the adoption of nanopores in applications outside of specialized academic labs. The DNA dose–response curve exhibits an elongated Gaussian shape—one similar to cell death response curves— that strongly indicate double–strand DNA breakage proceeds with three key characteristics: (i) indirect single– strand breakage by reaction with radicals, (ii) nearby SSBs on opposing strands generate a DSB, (iii) damage competes with bimolecular radical decay in the relevant dose range for clinical and emergency response applications. This agrees with prior results that suggest reaction with radicals, rather than direct damage with high energy particles, is the dominant mechanism of DNA damage in near physiological conditions.

Nanopore–based sensors thus show promise for a wide range of applications, including rapid dosimetry after accidental, imaging, or therapeutic radiation exposures. For instance, DNA damage due to different types of external beam radiation (x-rays, electrons, protons, etc.) or radiopharmaceuticals that emit many types of short-range radiation (e.g., soft x-rays, auger electrons, *α* particles, *β* particles), could be measured and characterized. We believe this data would provide quantitative information on the sensitivity of DNA to a range of different energies, linear energy transfer, dose rates and other properties of radiation. Such information would be a fundamental building block to new models and dose calculations that improve how we use radiation to treat cancer.

## IV. METHODS

### Sample preparation

2.5 kbp DNA fragments (Thermofisher, NoLimits) were exchanged from TE buffer (10 mmol*·*L^*−*1^ tris(hydroxymethyl)aminomethane, 2 mmol*·*L^*−*1^ ethylenediaminetetraacetic acid) to 20 mmol *·* L^*−*1^ sodium phosphate (pH = 7.4) using 1 kDa cut–off mini dialysis tubes (Cytiva) in order to minimize the effects of radical scavenging. The molecular concentration of DNA in the irradiated solution was 10 nmol · L^*−*1^. 5 kbp and 10 kbp DNA were added post irradiation as internal standards to calibrate for absolute DNA size and concentration. ^1^ This accounts for any variation in nanopipette geometry between runs. For irradiation, 300 *µ*L of solution was placed into eight 0.5 mL DNA LoBind tubes (Eppendorf, Cat. no. 022431005). Each solution was then exposed to a gamma–ray field from a Gammacell 220 ^60^Co irradiator (Nordion, Canada) for the time required to achieve the desired dose. The dose rate was 1.373 Gy/min and the irradiation temperature was 23 ^*◦*^ C. The uncertainty in absorbed dose in water is bounded by *±* 1.6 % for this irradiation geometry.

### Nanopipette measurement

Quartz nanopipettes were prepared with a laser assisted glass capillary puller (P-2000 Sutter Instruments, Novato, CA). Parameters such as the laser power, delay time, and strength of pull were adjusted to optimize the terminal diameter and interior geometry of the pipettes to maximize the measurement resolution of a particular analyte molecule or range of molecules. In this case, pipettes were optimized to achieve an acceptable success ratio in clearly defined log(ECD) frequency peaks for molecules in the 2.5 kbp to 10 kbp size range. The laser puller program used was as follows: HEAT = 575, VELOCITY = 25, DELAY = 180, PULL = 225. We use quartz capillaries with an external diameter of 1.0 mm and an internal diameter of 0.5 mm.

Nanopipettes were filled with a solution of 4 mol · L^*−*1^ LiCl and 10 mmol·L^*−*1^ TE buffer at pH 7.4. The exterior of the nanopipette was dipped into a 3 nmol · L^*−*1^ solution of the irradiated 2.5 kbp DNA, along with aliquots of the 5 kbp and 10 kbp internal standards (added at 0.3 nmol · L^*−*1^ concentration) in identical 4 mol · L^*−*1^ LiCl and 10 mmol·L^*−*1^ TE buffer. Each measurement was performed by a separate nanopipette to avoid cross contamination. Ionic current was monitored with an Axopatch 200B amplifier (Molecular Devices, San Jose, CA) sampling at 500 kHz, with a low pass bessel filter of 10 kHz. The ionic current data was analyzed using a threshold algorithm set at 5*σ*, where *σ* is the standard deviation of the ionic current to identify DNA-based resistive pulses. The signals were then decoded with a modified CUSUM-based algorithm [43] as implemented in the Nanolyzer software package (Northern Nanopore, Ottawa, Canada).

### Modeling

We consider a model for the DNA damage process that has two overall components. The first (combinatorial) component is how individually damaged nucleotides lead to double–strand DNA breaks (therefore reducing the intact DNA concentration we measure). The second is a reaction model for the damage of individual nucleotides. This reaction model provides the input—the defect probability, *P*_Λ_—into the first component. Otherwise, the two components are independent.

#### Component 1

When individual nucleotides are damaged along the double–helix, some fraction of those will result in a double–strand break. The damaged nucleotides—in particular, single–strand breaks—on opposing strands will likely lead to a double–strand break when they are within *l* − 1 base pairs of each other (i.e., for *l* = 1 this means that the two breaks have to be on a nucleotide pair). At room temperature and under normal salt conditions, *l* should be of the order of 5 base pairs. That is the scale when the base pairs in between two single–strand breaks can unravel and the double helix breaks into two fragment helices [42].

The model thus needs to find the likelihood of two defects being within *l* −1 base pairs of each other when every individual nucleotide has a probability *P*_Λ_ of being damaged. For a double–helix of length *L* base pairs, a configuration of damaged (“0”) and pristine (“1”) nucleotides will occur with probability

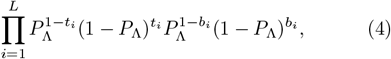

where *t*_*i*_ = 0 or 1 for the nucleotide on the top strand at position *i* and *b*_*i*_ = 0 or 1 for the bottom strand. Summing Eq. (4) over all configurations is equal to 1 and thus covers the configuration space.

Up to the resolution of detection, every original double–strand of DNA that has one or more double– strand breaks will reduce the intact DNA concentration. Thus, we need to sum Eq. (4) over all configurations that do not have any breaks, i.e., all configurations that do not have a damaged nucleotide on the top strand within *l* − 1 base pairs of a damaged nucleotide on the bottom strand. When *l* = 1 and only damaged nucleotides in a pair lead to a double–strand break, the sum should be over (*t*_*i*_, *b*_*i*_) = (1, 0), (0, 1), and (1, 1) for each *i*. This sum can be done sequentially, resulting in a total intact DNA helix probability, *P*, of

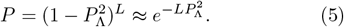

The approximate equality is from a Taylor expansion and is accurate for small *P*_Λ_. As a side note, 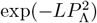 upper bounds 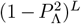 and this approximation can not explain the higher intact DNA concentrations that we observe at high doses.

Beyond this simple case, though, the sum over intact configurations will lead to a transfer matrix. However, the net effect of *l >* 1 is only to introduce an effective length scale 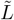, leading to

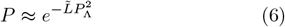

for small *P*_Λ_, which gives Eq. (1) when converted to a concentration. For the purposes here, this is all that is needed, since the fitting of experimental data employs aggregate parameters and thus cannot distinguish changes in 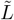 versus other parameters (see component 2 below).

In order to provide estimates of those other parameters, though, it is important to have a ballpark estimate of 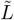. When *P*_Λ_ is small, the relationship of 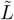 and *l* is as follows: When any given nucleotide on one strand is damaged (with probability *P*_Λ_), then the likelihood that there is damaged nucleotide on the opposing strand that will lead to a DSB is simply (2*l* − 1)*P*_Λ_. The overall probability, neglecting end effects, that there is damage on opposing strands within *l* of each other is then *LP*_Λ_ *·* (2*l −* 1)*P*_Λ_, giving an effective length of 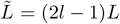. Considering *l ≈* 5, we expect 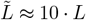, which is the value we use to estimate other fundamental parameters.

We note that the effective length absorbs not only the physical effect of how single–strand breaks lead to double–strand breaks but also measurement effects. When analyzing real data, 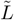 will also be impacted by measurement resolution. For instance, if the DNA breaks near its end, one will not be able to resolve the broken DNA from its intact form. Thus, one should subtract off some length (e.g., below about 100 base pairs for the measurements and analyte here). We work with a quite long sequence of DNA—2.5 kbp—and we do not expect the latter to be significant.

#### Component 2

The likelihood of a gamma photon directly interacting with a nucleotide is roughly in proportion to the ratio of the mass of DNA in solution to the mass of water (here, about 0.002 %). Therefore, under our conditions, radicals mediate almost all of the DNA damage. As the irradiated sample equilibrates, every •OH radical will react. ^*•*^OH can abstract hydrogen from the sugar of the nucleotide creating a SSB, or any of the bases which can propagate to a SSB or terminate in a host of non-scission terminating modifications [44]. All other hydroxyl radicals have to react with a radical scavenger, which includes other radicals produced upstream in radiolysis reactions [14].

Giving these considerations, we consider the reactions

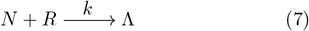

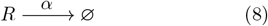

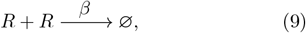

where *N* is the total nucleotide concentration, *R* the radical concentration, and Λ the total defect concentration. Defects occur via a bimolecular reaction of nucleotides and radicals with rate *k*. Radicals decay via a unimolecular process with rate *α* and a bimolecular process with rate *β*, both resulting in annihilation of the radicals (i.e., an inert chemical species, ∅). All reactions are taken to be irreversible since they all involve a substantial dissipation of energy. We emphasize that the unimolecular decay process does not influence the model fit to experimental data since it does not increase the number of effective (fit) parameters. However, it does influence the extraction of basic parameters and their interpretation. We also note that the reaction model, Eq. (7), does not consider any particular radical species or damage product. The underlying phenomenology is that the exact microscopic processes are unimportant to understand the emergence of double–strand breaks and the details are all absorbed by effective rate parameters.

The reactions above give rise to a set of rate equations,

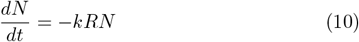

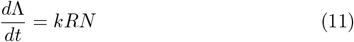

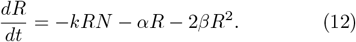

In addition to these rate equations, we need the initial concentrations. The initial total nucleotide concentration during irradiation is

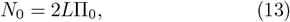

where *L* is the length of the DNA and Π_0_ is its initial concentration of DNA during irradiation (Π_0_ = 10 nM here). The initial radical concentration is taken to be linearly proportional to the radiation dose according to the well–known *G*–value [45],

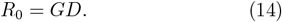

We assume that immediately after gamma irradiation the solution contains a uniform distribution of hydroxyl radicals and all nucleotides are equally likely to be damaged. Finally, the initial defect concentration is zero, Λ_0_ = 0.

When there are a lot of nucleotides compared to radicals (i.e., *N*_0_ ≫ *R*_0_) or the radical loss is fast compared to defect generation, then the depletion of nucleotides will be small, i.e., *N*_0_ ≫ Λ(*t → ∞*). Under these conditions, we can approximate the rate equations as

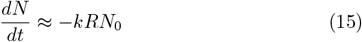

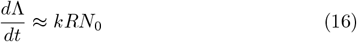

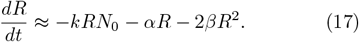

Replacing *N* with *N*_0_ on the right hand side of these equations will give a larger defect concentration at all times since *N* decreases monotonically with time. For the experimental results we present in this work, *P*_Λ_ *<* 0.01 for all conditions (i.e., all radiation doses). The magnitude of this probability indicates that replacing *N* with *N*_0_ is a good approximation. We note that *α* is an *effective* unimolecular decay parameter, since one decay channel is to react with nucleotides in a way that does not cause an SSB (i.e., *α* ∝ *N*_0_).

The approximate set of equations, Eq. (15) through Eq. (17), can be solved analytically. The solution for the radical concentration, taking *κ* = *kN*_0_ + *α*, is

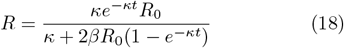

and the defect concentration is

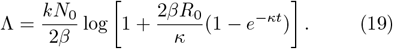

After all radicals react, the defect probability is

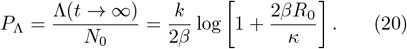

This is the main quantity of interest, Eq. (2), and is the input to the model of double–strand breaks, Eq. (1).

Finally, the fitting to Eq. (1) is done with the two effective fit parameters, Eq. (3), as we describe in the main text. Errors on the fit parameters are found via a boot– strapping calculation where the intact DNA concentrations are chosen according to a Gaussian distribution around their experimentally observed value with standard deviation given by the experimental error, and dose is taken to uniformly have a 1.6 % error. The prefactor— the intact DNA concentration, Φ_0_, when there is no irradiation—is taken as a known constant in each calculation within the boot–strapping process (i.e., Φ_0_ is taken as its value in the realization of noise). Errors for fundamental quantities come from the joint distribution of fit parameters.

## V. ACKNOWLEDGMENTS

We thank Mohamad Al-Sheikhly and Miral Dizdaroglu for sharing their deep understanding of the history of radiation chemistry and DNA.

M.L. acknowledges support from an NRC Postdoctoral Research Fellowship.

## Footnotes

1 Certain commercial materials, equipment, and instruments may be identified in this work to describe the experiments as completely as possible. In no case does such an identification imply a recommendation or endorsement by the National Institute of Standards and Technology, nor does it imply that the materials, equipment, or instrument identified are necessarily the best available for the purpose.

