## Supplemental Information for "Single–molecule biodosimetry"

In this supplemental information, we provide the details on the laser pulling process and settings, as well as details on the DNA concentration and raw nanopore signals.

### I. LASER PULLING PROCESS

We construct the pulled nanopipette pores as follows: We first clean 7.5 cm quartz capillaries (O.D. 1.0 mm, I.D. 0.50 mm) with an internal filament (Sutter instruments) by sonicating them upright in acetone for 15 minutes. They are then dried with compressed air and baked in a 70°C oven for 20 min. The pulling process uses a Sutter P-2000G laser–assisted pipette puller set to following parameters: HEAT = 575, FIL = 0, VEL = 25, DEL = 180, and PUL = 225. After pulling, we secure the nanopipettes horizontally and fill them with 4 M LiCl 10 mM TE buffer using 20 $\mu$L microloader pipette tips (Eppendorf). If an air bubble was visible near the nanopipette tip, then we refill the nanopipette in the same manner until the air bubble is removed.

### II. POST–IRRADIATION DNA CONCENTRATION

We prepare DNA solutions such that, after irradiation and dilution, the concentration of total nucleic acid is equivalent 3 nM (4.875 ng/$\mu$L) 2.5 kbp DNA. We confirm this after irradiation with UV-vis spectroscopy, Fig. S1.

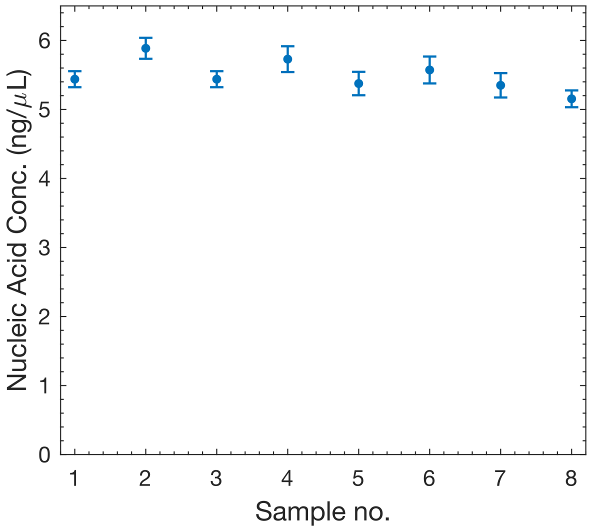


Fig. S1. **Post–irradiation DNA concentration.** The DNA concentration for each of the eight irradiated samples (at doses from 0 Gy to 15 Gy). We obtain the concentration with UV light absorbance using the NanoDrop One (Thermofisher). Error bars show plus/minus one standard deviation over 3 runs. The concentration is constant within the measurement error.

*

### III. IONIC CURRENT SIGNALS

Figure S2 shows the raw ionic current data (left panels) from each of the eight runs used to measure the intact DNA concentration as a function of dose, as well as direct uncorrected histograms (right panels). Each run begins with identically–prepared DNA solution from a single stock solution. As we stress in the main text, we collect each time–series with a unique nanopipette both to prevent cross contamination and to prevent fouling. We then use the Nanolyzer software package to extract and tabulate events (based on ECD). This data highlights a variety of different capture rates that appear uncorrelated to dose, as well as the variability in pore characteristics.

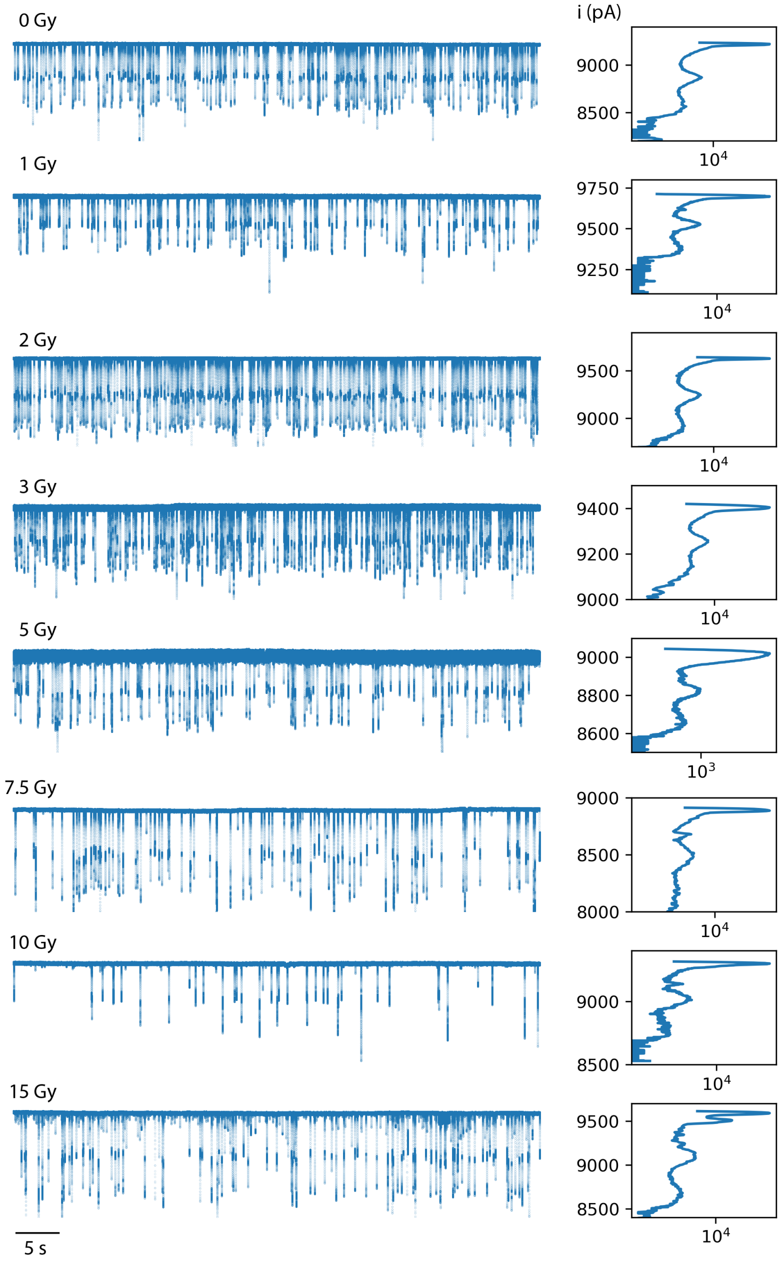


Fig. S2. **Ionic current time series.** From top to bottom, the left panels show the ionic current versus time for each of the doses. The right panels show the corresponding histogram for each of these real–time traces. This data highlights the variability of individual capillaries, from the baseline current to the positions of the different peaks. The dual internal molecular standards of this work correct for this variability and enable quantitative detection of nucleic acid analytes.
